# Challenges of Proper Disposal of Old Long-Lasting Insecticidal Nets and its Alternative Uses in Rural South-Eastern Tanzania

**DOI:** 10.1101/2022.12.01.518794

**Authors:** Sheila J. Salum, Winifrida P. Mponzi, Letus L. Muyaga, Joel D. Nkya, Yohana A. Mwalugelo, Marceline F. Finda, Hajirani M. Msuya, Dickson W. Lwetoijera, Emmanuel W. Kaindoa

## Abstract

**Introduction:** Insecticide-treated nets (ITNs) specifically long-lasting insecticidal nets (LLINs) are one of the most commonly used, scalable and cost-effective tools for controlling malaria transmission in sub-Saharan Africa. However, multiple alternative uses of retired LLINs have been observed and are associated with poor disposal practices. Nevertheless, the World Health Organisation (WHO) provided guidelines and recommendations for proper management of worn-out LLINs. This study assessed the existing alternative uses and disposal practices of old LLINs.

**Methods:** An explanatory sequential mixed-methods approach was used to assess LLINs existing alternative uses, disposal practices, knowledge and perceptions regarding WHO recommendations on proper disposal of old LLINs among stakeholders in Kilombero and Ulanga districts, southe-astern Tanzania. A survey questionnaire was administered to 384 respondents, Focus Group Discussions (FGD) and Key Informant Interviews (KII) were conducted to clarify responses regarding existing disposal practices with associated challenges and alternative uses of the LLINs. Findings from both study components were used to draw inferences.

**Results:** A total of 384 people surveyed, 97% were not aware of the WHO recommendation on proper disposal of old LLINs. The common methods used to dispose LLINs were burning 30.73%, disposing of into garbage pit 14.84% and alternative uses 12.24 %. Of respondents with LLINs (239); 41% had alternative uses while 59% had no alternative uses. The common alternative uses were ropes for tying or covering items 20.92%, garden fencing 7.53%, chicken coops 5.02% and 7.53% for other minor alternative uses. All key informants reported being unaware of the WHO guideline on the proper disposal of the old LLINs.

**Conclusion:** This study demonstrates that despite participant’s limited knowledge on WHO guidelines for proper disposal of old LLINs, after presenting these guidelines, majority are willing to comply. Comprehensive efforts are therefore needed to address challenges associated with poor disposal, alternative uses and awareness about WHO guidelines among key stakeholders. Collection strategies should be agreed upon within the community members prior to replacement. Since alternative uses sometimes referred to as repurposing of old nets, proper guidelines should be developed to ensure that repurposing of old LLINs do not cause harm to human health and the environment.

## Background

There is a growing environmental concern attributed to poor disposal of old Insecticide-Treated Nets (ITNs) specifically long-lasting insecticidal nets (LLINs), which necessitate urgent actions. Globally more than 2.3 billion ITNs had been distributed between 2004–2020 period, and sub-Saharan Africa (SSA) received 2 billion (86%) of all distributed insecticide-treated nets [1]. In 2020, more than 230 million of ITNs had been distributed globally by National Malaria Control Programmes (NMCP) in malaria endemic countries and more than 194 million were distributed in SSA, and Tanzania was among the top five recipients [1]. This massive volume of LLINs placed in circulation could lead to the rapid increase of solid waste litter and environmental pollution in SSA countries where solid waste management is still a major challenge [2,3].

The primary aim of using LLIN is to protect people against mosquito bites, malaria transmission and other mosquito borne infections [4]. Though, in several parts of Africa, LLINs have been reported to be used for fencing vegetable gardens and chicken huts, sieving grains, and as fishing nets and alternative building materials, which may lead to environmental harm [5–9]. However, the insecticides (pyrethroids and pyrroles) embedded in the LLINs though have low mammalian toxicity, if not properly disposed, it may be toxic to aquatic life and other untargeted species. Worryingly, these insecticides might exert additional insecticide selection pressure on mosquitoes at their aquatic habitats [10]. Thus, any forms of repurposing should consider protecting such life as stipulated in the sustainable development goals (SDGs) 14 and 15 on Life below water and Life on Land respectively [11].

Plastic pollution is among the public health concerns due to its abundance, durability, and persistence in the environment as well as the perceived threats to living organisms [12]. Environmental pollution resulting from plastic materials continues to be a global challenge due to threats to aquatic life, wildlife, climate change, human health, and economic development [13–17]. The current evidence indicates that manufactured plastic materials have increased annually, thus environmental hazard has also significantly increased and become a global concern [18]. Globally, about 10% of the total municipal waste is plastic constituting over 90% of the ocean debris [13,17]. The estimates show that about1600 million tonnes of plastic wastes will be produced in 2050 unless measures are taken to limit the production [19]. Being among the possible sources of plastic waste, up to date, more than 2.3 billion long-lasting insecticidal nets (LLINs) have been distributed globally [1] whose fate is unknown. Despite the adoption of an anti-plastic bag policy in various countries in SSA [20], there have been deficient measures against LLINs and their packages/bags disposal. The WHO has provided warnings on the possible impacts of accumulated LLINs and their packages, which may include: contamination of crops/vegetables, soil, and groundwater when these LLINs are reused and may also produce dangerous persistent toxins resulting from open-air burning [21]. Thus, the development of environmentally friendly programs for proper disposal of LLINs and their packing bags should be explored to adhere to the current legislation specified by environmental management authorities. Nevertheless, World Health Organization (WHO) has recommended the proper ways of disposing or handling the old LLINs such as; 1) continue to be used even if they have holes until they are replaced with new LLINs, 2) not to dispose of the old LLINs in water bodies, 3) collection of the old LLINs for disposal by National Malaria programs, 4) incineration of old LLINs 5) formulating guidelines, policies, and regulations by the NMCPs in collaboration with national environmental authorities [21,22]. An important question is whether local health and environmental authorities are aware of such guidelines and whether they have capacity to implement and monitor these guidelines. Yet, the evidence from multiple sources indicates high levels of ITNs attrition from households [23], indicating high concerns of accumulation of old nets in the environment.

Despite clear information from relevant authorities including; WHO, United Nations Environment Programme (UNEP), Basel, Rotterdam and Stockholm Conventions on sound management of old LLINs, there is evidence of misuse and improper disposal of the old LLINs. Evidence show that the communities still use LLINs for other activities rather than using for protection against mosquito biting [7]. Thus, this study was conducted to assess existing alternative uses and disposal practices of the LLINs as well as community members and key stakeholders’ knowledge and perception regarding the WHO recommendations on proper disposal of LLINs in Kilombero valley, south-eastern Tanzania. Therefore, understanding the factors influencing poor disposal practices and associated alternative uses would be beneficial for decision and policy makers.

## Methods

### Study area

This study was conducted in six wards in Kilombero and Ulanga districts, both in the Kilombero valley in south-eastern Tanzania (Figure 1). In Ulanga district, the study was done in Kivukoni, Mavimba and Minepa villages. In urban settings of the Kilombero district, the study was done in Ifakara town council and its surrounding wards including Katindiuka, Mlabani and Mbasa. The main economic activities in the study sites are predominantly rice farming, retail businesses, and fishing. The major mosquito control intervention is use of LLINs that are freely distributed at antenatal clinics and School Net Program (SNP) and universally distributed by the government through every 3–4 years [24] and the last mass distribution by the government was done in 2016.

**Figure 1:**
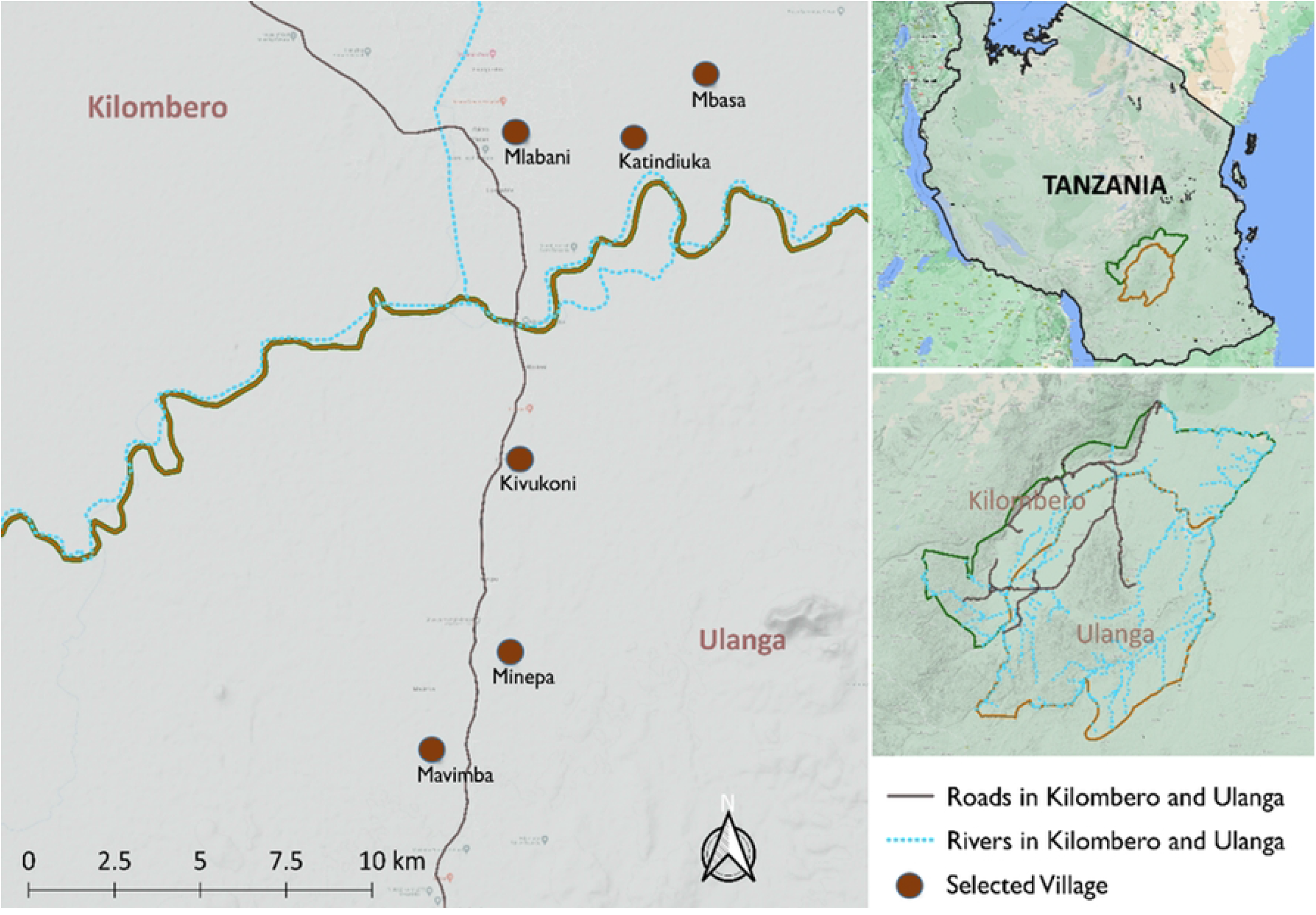
A map of Kilombero and Ulanga districts in south-eastern Tanzania showing study wards.

## Study design and data collection

This study was an explanatory sequential mixed-method design [25,26]; quantitative analysis data provided insights for qualitative study themes. This approach had two components: the first arm was a quantitative survey of 384 households in two districts. The second arm consisted of four focus group discussions (FGDs) with community members selected from survey respondents (two groups with males and two with females) followed by four key informants from public health and environmental sectors. The qualitative findings were used to clarify some of the responses from the initial survey questionnaire (Figure 2). Findings from the two components were used to make inferences.

**Figure 2:**
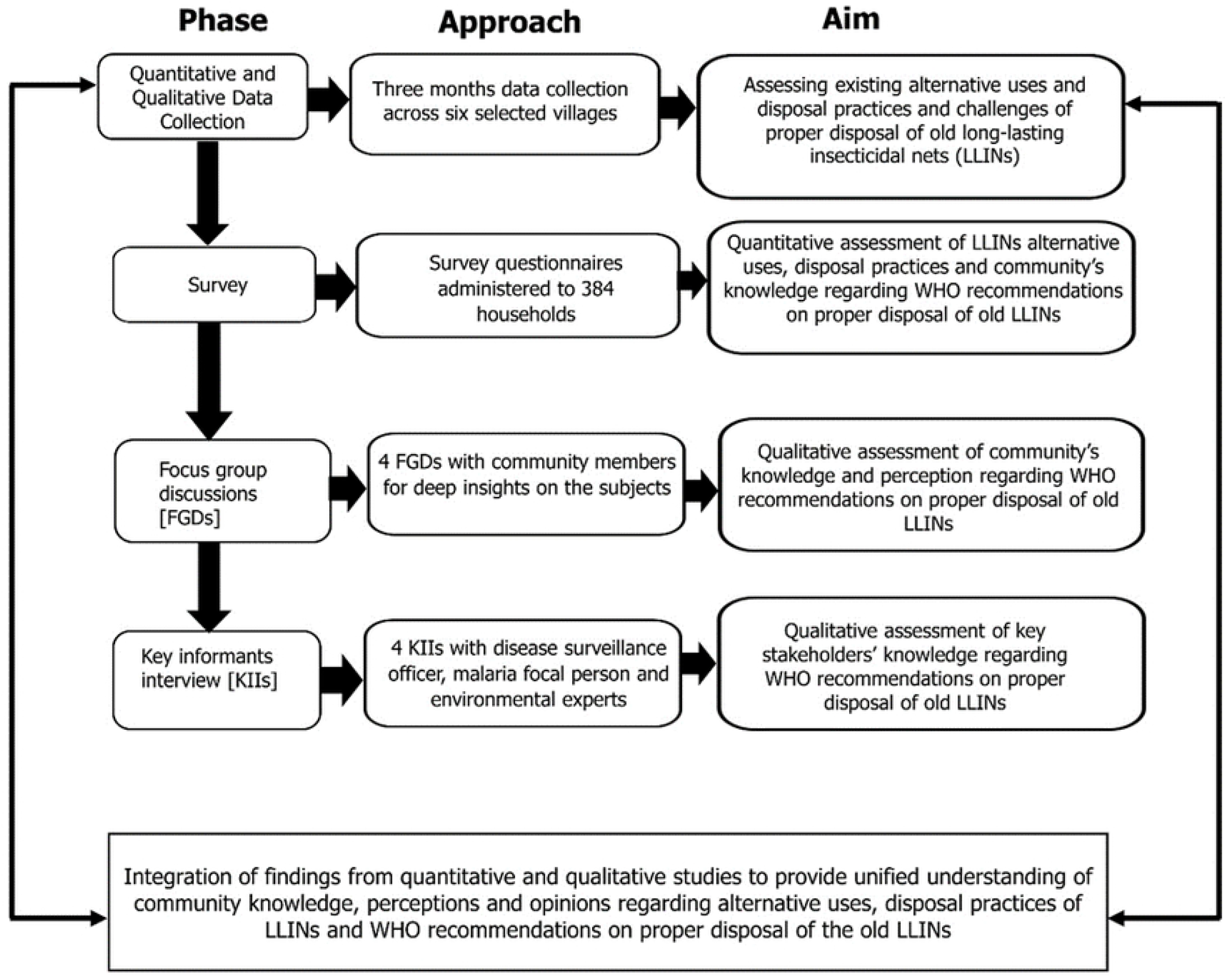
Illustration of the explanatory sequential mixed methods approach used to examine community knowledge and perceptions of alternative uses, disposal practices and challenges and WHO recommendations regarding proper disposal of old LLINs.

## Survey questionnaire

The sample size for the survey questionnaire was determined using Epi-Info Software package [27] based on the following assumptions: sample size based on the single proportion for cross-sectional survey assumed; 50% of the population would be observed with poor disposal and alternative uses of bed nets [28,29], with 5% margin of error at 95% confidence level. Thus, a sample size of 384 was reached, however sample size was approximated to 400 to make room for the non-response at 5% rate. To ensure balanced design, 64 households were randomly selected on each ward and consented to participate in the survey. A survey questionnaire was translated in Swahili and administered using electronic forms on a free-access application programmed on Open Data Kit (ODK) from KoboTool Box server [30]. Later the survey was then translated back from Swahili into English. The questionnaire was administered to 10 different potential participants for piloting and was revised accordingly.

## Focus group discussions

Four (4) focus group discussions (FGDs) were held in May 2022 with a subset of survey respondents to clarify the quantitative component’s findings. Participants were purposively selected; they included those who had or did not have long-lasting insecticidal nets (LLINs), those who had alternatively used and those who had not alternatively used the bed nets. The discussions consisted of two male groups and two female groups in four wards; each session had 7-8 participants. To maximize participation, male and female participants were separated during the discussions. The discussions provided detailed information on community members’ understanding and perceptions of alternative uses of LLINs, LLINs disposal practices and awareness and understanding of WHO guideline on proper management of old LLINs. The topics explored in the FGDs included experiences and perceptions of the participants on 1) LLIN use, care and repair; 2) LLIN damage and changing decision; 3) LLINs disposal of and alternative uses and; 4) WHO guideline awareness on the sound management of old LLINs. The FGD guide was pre-tested and questions were refined according to the outcomes, the guide was then translated from Swahili into English. The discussions were held at local ward offices or in classrooms at local primary schools. The discussions were audio-recorded and held in Swahili, Tanzania’s native language.

## Key Informant Interviews

Key informant interviews (KIIs) were conducted with four purposively selected stakeholders from public health and environmental sectors, who had a direct or indirect involvement in malaria, vector control and environmental management. The KIIs included Ifakara disease surveillance officer, Ulanga malaria focal person, Ifakara environmental officer, Ulanga environmental health officer. These informants were interviewed to investigate their knowledge regarding WHO recommendations as well as their awareness of ongoing bed net alternative uses and disposal practices. The discussions provided in-depth information on key stakeholders’ perceptions and perspectives on 1) malaria vector control interventions, progress and challenges; 2) alternative uses and disposal challenges of the old LLINs 3) WHO guideline awareness on proper management of LLINs. The interviews were conducted in Swahili language and audio-recorded, specific notes were taken.

## Data processing and analysis

All survey data were extracted from KoboTool Box software [30], then checked and cleaned in Excel, and finally coded in R statistical software version 4.1.2 [31]. The data analysis was mainly descriptive, and statistics were presented as mean and standard deviations (SD) for continuous variables, and categorical variables were summarized using frequencies and percentages. The analysis involved some inferential statistics, i.e., Chi square and logistic regression. A Pearson’s Chi-square test was performed to understand the factors that were associated with the alternative uses and disposal practices. Bivariate analysis was done using binary logistic regression model whereby *glm ()* function with *logit* link was used to assess the influence of socio-demographic characteristics and other independent variables related to net ownership on alternative use of old LLINs. Additionally, stepwise logistic regression was adopted to obtain the best performing model using *stepAIC ()* function from MASS package.

Audio data from FGDs and KIIs were transcribed and then translated from Swahili to English. Notes taken during the discussions were incorporated into the written transcripts. The transcripts were then imported into Atlas.ti.22 software [32] for coding. Survey findings informed development of KII and FGD guides, and then the guides informed deductive coding.

Findings were presented using the integration principles and practices in mixed methods designs as described by Fetters *et al*., [33]. Weaving approach was used, in which both qualitative and quantitative findings were reported together based on the illustrated themes in Figure 2. Quantitative findings from the survey were presented, and explanations for some of the concepts were given from the FGDs. Direct quotations from the FGD participants were reported in some selected cases to further describe the themes.

## Ethical considerations

Ethical approvals to conduct this study were provided by the Medical Research Coordinating Committee of the National Institute for Medical Research of Tanzania (NIMR) with approval number (Ref: NIMR/HQ/R.8a/Vol. IX/3353) and Institution Review Board (IRB) of Ifakara Health Institute, approval number IHI/IRB/No:12-2022. Before conducting the study, the District Medical Officers were informed about the study and granted permission and informed all local leaders; through an introduction letters Ref No. IHI/ADM/22/0689 and IHI/ADM/22/0690. Additionally, consent to conduct the study was sought from both, communal and individual level. Communal consent was from face-to-face discussion with local leaders about the study and request to conduct it in their wards. Individual consent was by discussing with each participant about the study procedures and its implications/importance followed by a request to participate. Those who agreed to participate were given a written consent forms to fill before filling in the questionnaires. Permission to publish this manuscript was obtained from National Institute of Medical Research (Ref: NIMR/HQ/P.12 VOL XXXV/77).

## Results

### Demographic characteristics of survey respondents

A total of 384 household representatives (192 from peri-urban and 192 from rural) responded to the survey questionnaire of which 67.5% were females and 32.5% males. With regard to occupation status; 359 were farmers, less than 12% engaged in business activities and fewer (8%) were engaged in other activities. More than three-quarter of the participants (83.3%) had a primary education, 16.7% had a secondary and above. Nearly two-thirds (67.7%) of the participants were married or living with a partner, 23.3% were not married, and 9.1% were widowed. More than 50% of the interviewed households had household size of 4 to 6 people (Table 1).

**Table 1:**
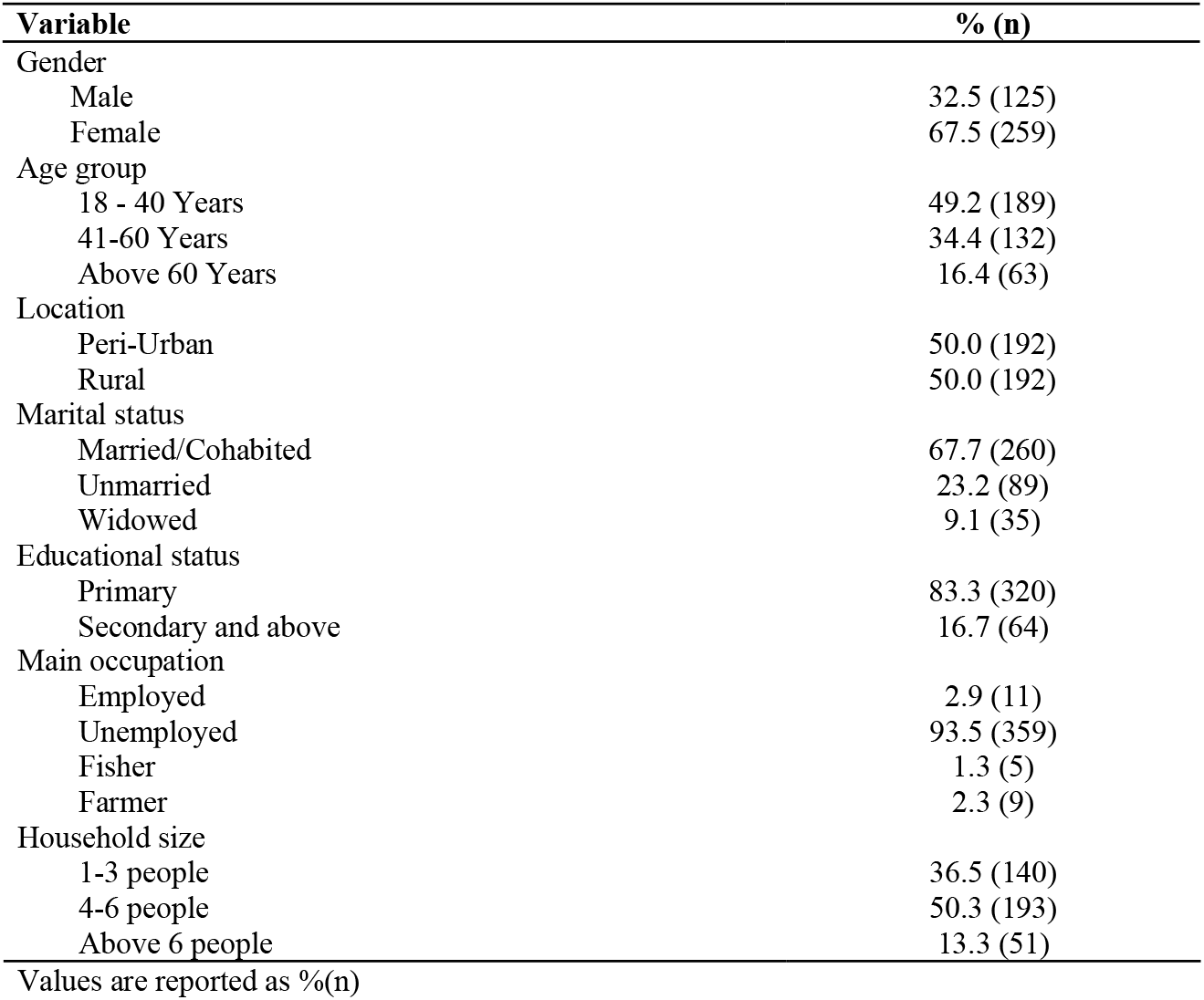
Demographic information of the study participants (N = 384)

| Variable | % (n) |
| --- | --- |
| Gender |  |
| Male | 32.5 (125) |
| Female | 67.5 (259) |
| Age group |  |
| 18 - 40 Years | 49.2 (189) |
| 41-60 Years | 34.4 (132) |
| Above 60 Years | 16.4 (63) |
| Location |  |
| Peri-Urban | 50.0 (192) |
| Rural | 50.0 (192) |
| Marital status |  |
| Married/Cohabited | 67.7 (260) |
| Unmarried | 23.2 (89) |
| Widowed | 9.1 (35) |
| Educational status |  |
| Primary | 83.3 (320) |
| Secondary and above | 16.7 (64) |
| Main occupation |  |
| Employed | 2.9 (11) |
| Unemployed | 93.5 (359) |
| Fisher | 1.3 (5) |
| Farmer | 2.3 (9) |
| Household size |  |
| 1-3 people | 36.5 (140) |
| 4-6 people | 50.3 (193) |
| Above 6 people | 13.3 (51) |

Values are reported as %(n)

### Alternative uses from of survey respondents

Chi-square test of proportion was used to associate alternative uses with social demographic characteristics of the study participants, LLINs sources, disposal determinants and WHO guideline awareness as presented on Table 3. Of 239 study participants with LLINs, 98 had LLINs alternative use. About 69.4% of the study participants who had LLINs alternative use were female. Participants who reported to have alternative use of LLINs (56.1%) aged between 18 to 40 years. More than half (51.2%) of the participants with LLINs alternative uses were from rural setting. With regard to marital status, about 63.3% of participants were married or cohabiting. Majority of participants who reported to have alternative use of LLINs (81.6%) had primary education while 18.4% had secondary education. More than 90% of the participants who practiced alternative uses were farmers.

Most of the participants who had more than two LLINs 74 (75.5%) were found to have more alternative uses compared to those with less than two LLINs 24 (24.5%) (p-value < 0.001). Bed nets donation sources were mainly obtained from SNP 28 (28.6%), clinic 16 (16.3%), mass distribution 6 (6.1%) and fewer from colleagues 2 (2.0%), with (p-value < 0.001) (Table 3). Majority of the LLINs used for alternatives uses were mainly used by children aged less than 5 years 59 (60.2%), children aged 6 −17 years 24 (24.5%), adults aged 18-50 years 12 (12.2%) and fewer from elders aged above 50 years 3 (3.1%); (p-value < 0.001). The participants were also asked on how often they change their LLINs as it might influence alternative use. Interestingly, more than 50% of the respondents reported to change their LLINs within 6 months to 2 years; p-value < 0.001. Furthermore, the changing decision of the old or worn-out LLINs were 74 (75.5%), torn 19 (19.4%) and fewer were other reasons 5 (5.1%) with p-value < 0.001 (Table 2). Other variables such as age, gender, marital status, education status, location, occupation and household size were not statistically significant associated with the alternative uses of LLINs.

**Table 2:**
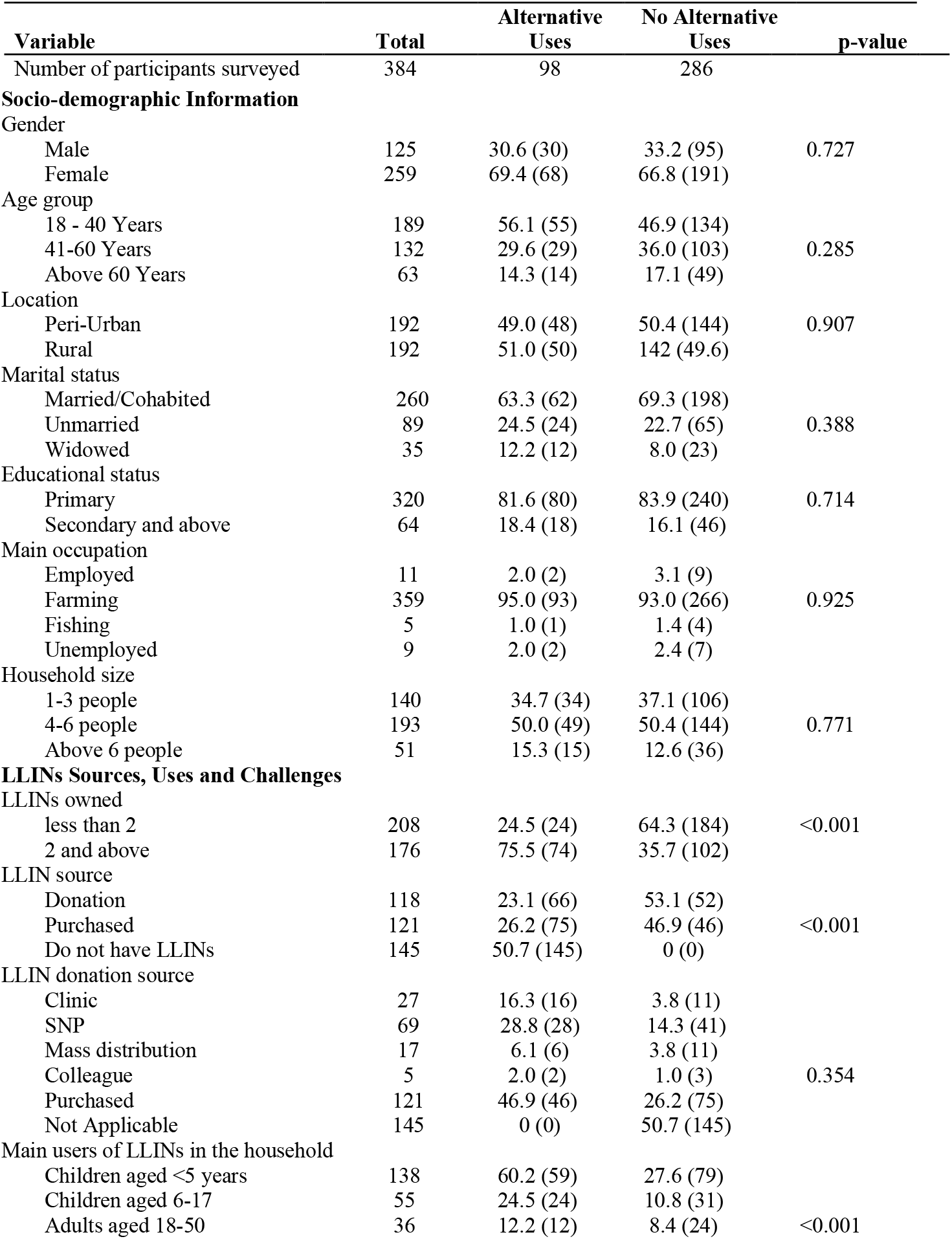
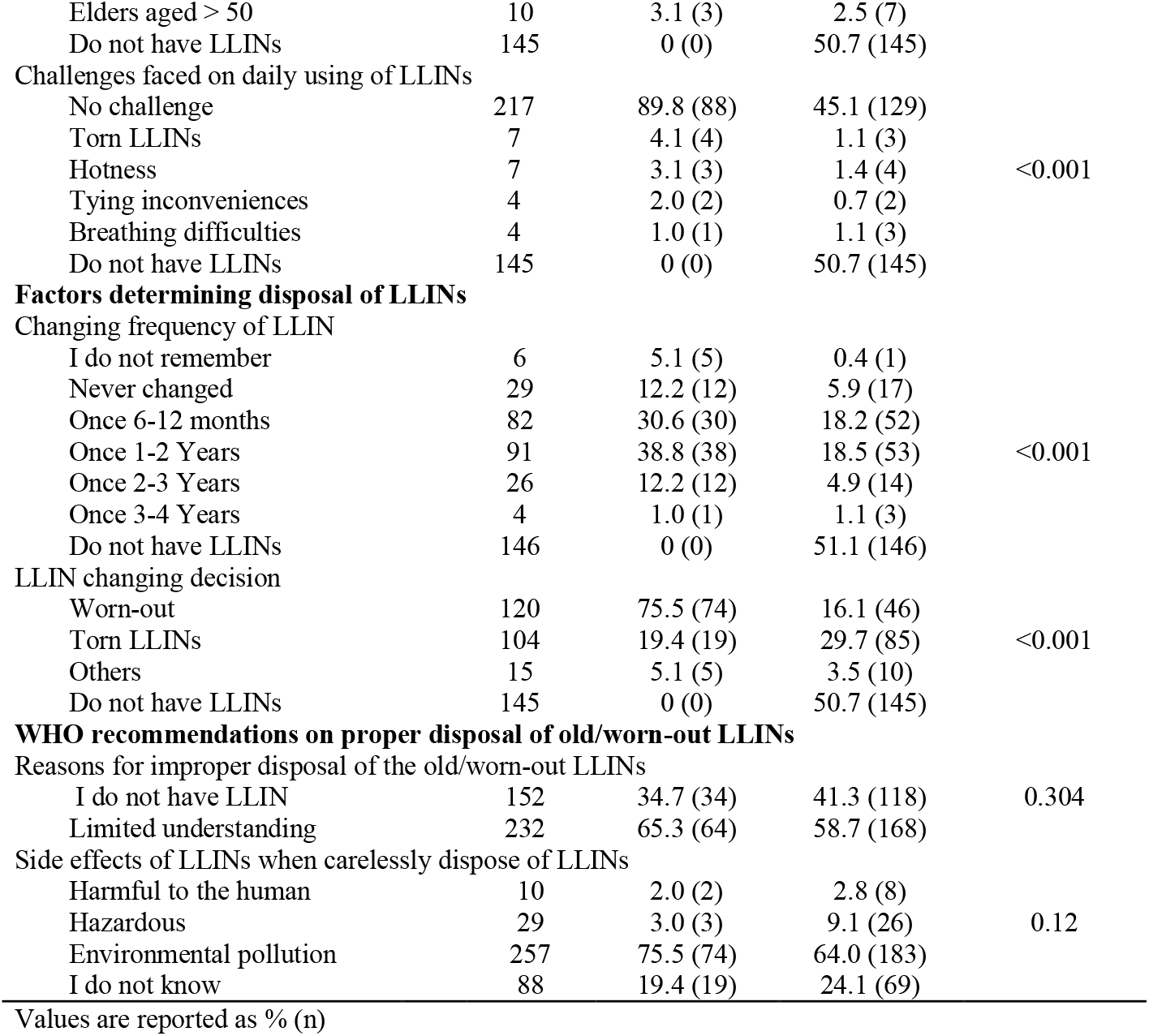
Alternative use practices and disposal challenges of LLINs (N = 384)

| Variable | Total | Alternative Uses | No Alternative Uses | p-value |
| --- | --- | --- | --- | --- |
| Number of participants surveyed | 384 | 98 | 286 |  |
| Socio-demographic Information |  |  |  |  |
| Gender |  |  |  |  |
| Male | 125 | 30.6 (30) | 33.2 (95) | 0.727 |
| Female | 259 | 69.4 (68) | 66.8 (191) |  |
| Age group |  |  |  |  |
| 18 - 40 Years | 189 | 56.1 (55) | 46.9 (134) |  |
| 41-60 Years | 132 | 29.6 (29) | 36.0 (103) | 0.285 |
| Above 60 Years | 63 | 14.3 (14) | 17.1 (49) |  |
| Location |  |  |  |  |
| Peri-Urban | 192 | 49.0 (48) | 50.4 (144) | 0.907 |
| Rural | 192 | 51.0 (50) | 142 (49.6) |  |
| Marital status |  |  |  |  |
| Married/Cohabited | 260 | 63.3 (62) | 69.3 (198) |  |
| Unmarried | 89 | 24.5 (24) | 22.7 (65) | 0.388 |
| Widowed | 35 | 12.2 (12) | 8.0 (23) |  |
| Educational status |  |  |  |  |
| Primary | 320 | 81.6 (80) | 83.9 (240) | 0.714 |
| Secondary and above | 64 | 18.4 (18) | 16.1 (46) |  |
| Main occupation |  |  |  |  |
| Employed | 11 | 2.0 (2) | 3.1 (9) |  |
| Farming | 359 | 95.0 (93) | 93.0 (266) | 0.925 |
| Fishing | 5 | 1.0 (1) | 1.4 (4) |  |
| Unemployed | 9 | 2.0 (2) | 2.4 (7) |  |
| Household size |  |  |  |  |
| 1-3 people | 140 | 34.7 (34) | 37.1 (106) |  |
| 4-6 people | 193 | 50.0 (49) | 50.4 (144) | 0.771 |
| Above 6 people | 51 | 15.3 (15) | 12.6 (36) |  |
| **LLINs Sources, Uses and Challenges** |  |  |  |  |
| LLINs owned |  |  |  |  |
| less than 2 | 208 | 24.5 (24) | 64.3 (184) | <0.001 |
| 2 and above | 176 | 75.5 (74) | 35.7 (102) |  |
| LLIN source |  |  |  |  |
| Donation | 118 | 23.1 (66) | 53.1 (52) |  |
| Purchased | 121 | 26.2 (75) | 46.9 (46) | <0.001 |
| Do not have LLINs | 145 | 50.7 (145) | 0 (0) |  |
| LLIN donation source |  |  |  |  |
| Clinic | 27 | 16.3 (16) | 3.8 (11) |  |
| SNP | 69 | 28.8 (28) | 14.3 (41) |  |
| Mass distribution | 17 | 6.1 (6) | 3.8 (11) |  |
| Colleague | 5 | 2.0 (2) | 1.0 (3) | 0.354 |
| Purchased | 121 | 46.9 (46) | 26.2 (75) |  |
| Not Applicable | 145 | 0 (0) | 50.7 (145) |  |
| Main users of LLINs in the household |  |  |  |  |
| Children aged <5 years | 138 | 60.2 (59) | 27.6 (79) |  |
| Children aged 6-17 | 55 | 24.5 (24) | 10.8 (31) |  |
| Adults aged 18-50 | 36 | 12.2 (12) | 8.4 (24) | <0.001 |
| Elders aged > 50 | 10 | 3.1 (3) | 2.5 (7) |  |
| Do not have LLINs | 145 | 0 (0) | 50.7 (145) |  |
| Challenges faced on daily using of LLINs |  |  |  |  |
| No challenge | 217 | 89.8 (88) | 45.1 (129) |  |
| Torn LLINs | 7 | 4.1 (4) | 1.1 (3) |  |
| Hotness | 7 | 3.1 (3) | 1.4 (4) | <0.001 |
| Tying inconveniences | 4 | 2.0 (2) | 0.7 (2) |  |
| Breathing difficulties | 4 | 1.0 (1) | 1.1 (3) |  |
| Do not have LLINs | 145 | 0 (0) | 50.7 (145) |  |
| **Factors determining disposal of LLINs** |  |  |  |  |
| Changing frequency of LLIN |  |  |  |  |
| I do not remember | 6 | 5.1 (5) | 0.4 (1) |  |
| Never changed | 29 | 12.2 (12) | 5.9 (17) |  |
| Once 6-12 months | 82 | 30.6 (30) | 18.2 (52) |  |
| Once 1-2 Years | 91 | 38.8 (38) | 18.5 (53) | <0.001 |
| Once 2-3 Years | 26 | 12.2 (12) | 4.9 (14) |  |
| Once 3-4 Years | 4 | 1.0 (1) | 1.1 (3) |  |
| Do not have LLINs | 146 | 0 (0) | 51.1 (146) |  |
| LLIN changing decision |  |  |  |  |
| Worn-out | 120 | 75.5 (74) | 16.1 (46) |  |
| Torn LLINs | 104 | 19.4 (19) | 29.7 (85) | <0.001 |
| Others | 15 | 5.1 (5) | 3.5 (10) |  |
| Do not have LLINs | 145 | 0 (0) | 50.7 (145) |  |
| **WHO recommendations on proper disposal of old/worn-out LLINs** |  |  |  |  |
| Reasons for improper disposal of the old/worn-out LLINs |  |  |  |  |
| I do not have LLIN | 152 | 34.7 (34) | 41.3 (118) | 0.304 |
| Limited understanding | 232 | 65.3 (64) | 58.7 (168) |  |
| Side effects of LLINs when carelessly dispose of LLINs |  |  |  |  |
| Harmful to the human | 10 | 2.0 (2) | 2.8 (8) |  |
| Hazardous | 29 | 3.0 (3) | 9.1 (26) | 0.12 |
| Environmental pollution | 257 | 75.5 (74) | 64.0 (183) |  |
| I do not know | 88 | 19.4 (19) | 24.1 (69) |  |

Values are reported as % (n)

In addition, logistics regression was fitted to access the strength and magnitude of association between alternative uses and others predictor variables. The results from fitted logistic regression indicated that LLINs donation sources and changing decision were significantly associated with alternative use. After adjusting for other factors, participants who obtain the LLINs from clinic had about 5-fold higher probability of having alternative use of LLINs as compared to participants who obtain from mass distributions (OR= 05.367, 95% CI: 0.472-6.397) (Table 2). Interestingly, the changing decision of LLINs due to being torn was statistically significant associated with 90%less odds (OR=0.113, 95% CI: 1.224-23.540) of having alternative use as compared to when the net is worn-out (Table 3).

**Table 3:**
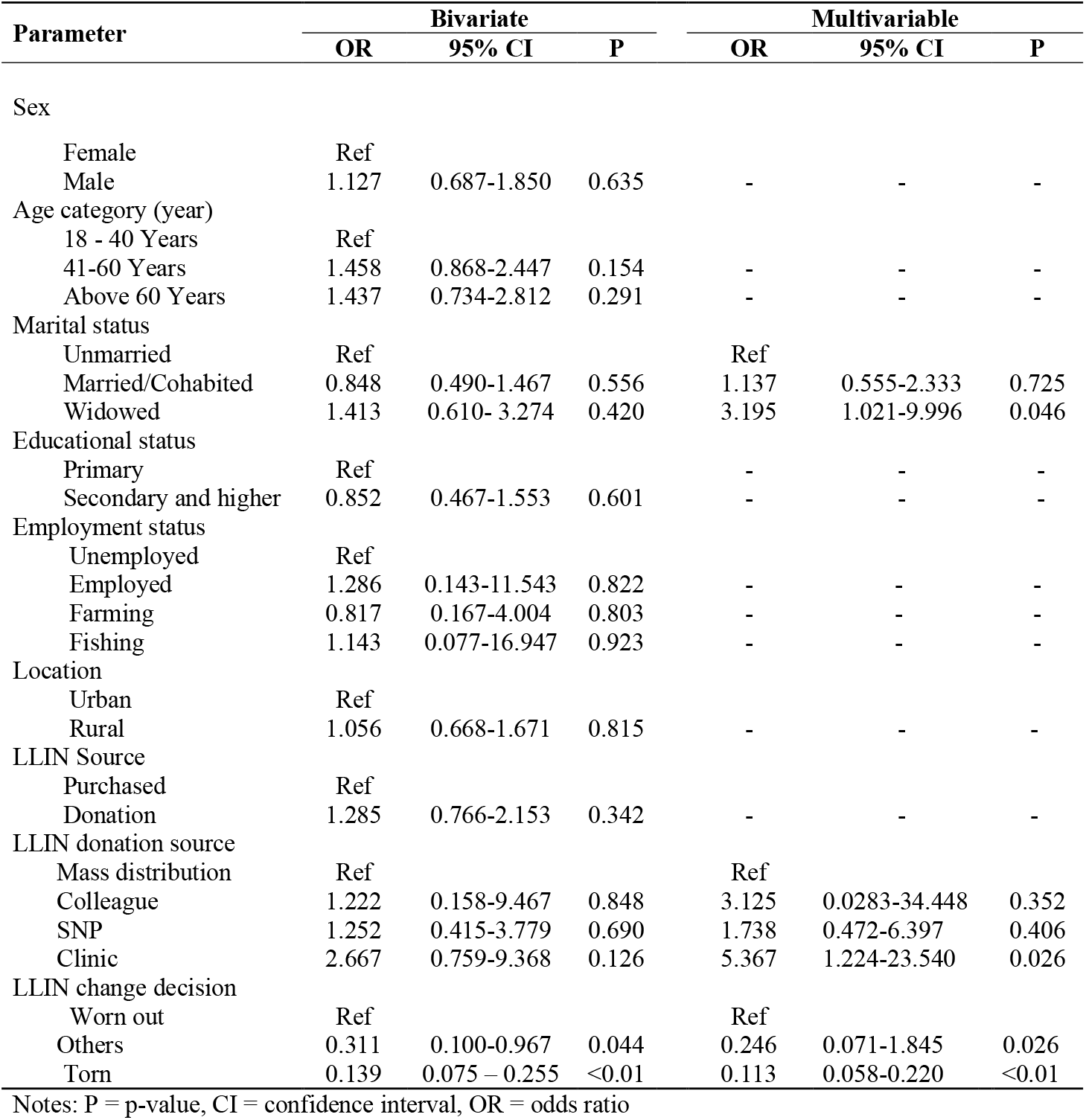
Predictors of Alternative use practices and disposal of LLINs (N = 384)

## Primary themes generated

Four (4) themes emerged from the analysis of the FGDs and KIIs scripts: 1) general knowledge on malaria prevention measures, 2) what happens past useful phase, 3) awareness and perceptions of WHO guideline, and 4) recommendations for future for improving disposal mechanisms of LLINs.

### General knowledge on malaria prevention measures

Almost two thirds of the respondents 62.2% (n=384) had bed nets particularly long-lasting insecticidal nets (LLINs) and use them on regular basis. Respondents were aware of malaria transmission and appropriate protective measures, due to extensive awareness campaigns raised from the Tanzania Ministry of Health and Non-governmental organizations. The most commonly reported prevention measures were LLINs, mosquito coils, skin mosquito repellents, long clothes, larval source management (LSM) and aerosol space sprays. Other mentioned preventive tools were smoky fires from specific leaves or barks and burning of unused clothes to chase away mosquitoes for some time before bed time. The community members mostly used bed nets while indoors with minimal protection while outdoors as this community member said:

> *“First of all, I always pay attention to the bed net tuck-in time thus by 06:00 PM in the evening to avoid mosquitoes from entering into the bed net. While we are outside, we do not use any special tool to protect ourselves, we only wave to chase away the mosquitoes though they keep on biting us. When we are done with the outdoor chores/activities, we go inside and sleep under LLIN believing there are no mosquitoes” (Male, Kivukoni).*

### Perceptions of community members on bed nets uses, effectiveness and challenges

Bed nets were reported to be a very effective malaria control method, the majority of FGD participants had positive attitude toward bed nets use, though they expressed frustration that malaria remained a problem despite regular use of LLINs. Participants’ responses to the effectiveness of LLINs in malaria prevention was more salient subjective factors. They claimed that effectiveness of LLINs was inhibited or reduced in several circumstances, including staying outside during evening and the need to run chores which may result in outdoor mosquito bites as this community member reported:

> *“In my understanding, the use of LLIN can be effective however it depends under what circumstance, for example staying outdoor environment in the evening while running chores one can be bitten by mosquitoes before entering in the bed net” (Female, Katindiuka).*

Furthermore, some participants argued that depending on LLIN only is not the solution to prevent one from getting malaria if LLIN is not well cared and repaired, mosquito breeding sites are not eradicated, doors and windows are left opened until sunset.

> *“On my opinion, LLIN is not effective way to prevent malaria because if you only use LLIN while your outdoor environment is not clean, mosquito breeding sites are not cleared therefore even though you sleep under LLIN you do not prevent malaria” (Female, Katindiuka).*

A majority of the participants had positive attitude towards sleeping under LLINs to avoid mosquito bites and malaria infection. Although there were some challenges to using bed nets on a daily basis, particularly LLINs, these include LLIN durability, large mesh and being small, and an irritating odor when the LLIN is new triggering cough like reaction. Participants expressed dissatisfaction with some LLINs due to unfriendly characteristics such as its large mesh size that is perceived to allow small mosquitoes to penetrate through. Further, when repaired, LLINs tend to shrink and shorten, making them difficult to keep securely tucked in under sleeping beds/mattresses.

> *“The donated LLINs have large mesh/holes to the extent that the tiny mosquito can penetrate through believing you have protected yourself from mosquito bites by the time you realize the mosquito has already suck blood from you. So, you start asking yourself when did it enter as before entering in the LLIN you made sure it is well tucked in and chased all the mosquitoes inside” (Female, Minepa).*

### Bed nets changing habit

Majority of the participants reported to change their bed nets every after 6 months to 3 years depending on tearing. The lifespan of LLINs for mosquito protection is locally perceived to decline in the first 6 months with the visible holes or tears. Participants’ responses to the reasons for bed net change were similar across all focus groups, the major reasons were LLIN having many holes, torn or worn-out as it is perceived fading of the insecticide, making the bed net unfit for the protection purpose (Table 4). Although, most of the FGD participants stated that age of the LLIN was not the reason for them to change their nets as long as it was still in a good condition (referring to the number of holes), repairs made on the nets and absence of bed net to replace due to economical reason.

> *“I change the LLIN twice in a year, the reasons for changing the LLIN is worn-out or fading of the insecticide when I see mosquitoes inside the LLIN then I am forced to change the LLIN” (Female, Minepa).*

**Table 4:** Malaria prevention methods, nets changing habit, existing LLINs alternative use practices and their causes

| Variable assessed | Participant responses | % (n) |
| --- | --- | --- |
| LLINs ownership (n=384) | Yes | 62.24 (239) |
|  | No | 37.76 (145) |
| Number of LLINs owned (n=384) | <2 | 54.17 (208) |
|  | ≥2 | 45.83 (176) |
| Bed nets changing frequency (n=239) | Once 6-12 months | 34.31 (82) |
|  | Once 1-2 years | 38.07 (91) |
|  | Once >2 years | 13 (31) |
|  | Forgot | 2.51 (6) |
|  | Never changed | 12.13 (29) |
| Bed nets changing decision (n=239) | LLINs being worn-out | 43.51 (104) |
|  | LLINs being torn | 50.21 (120) |
|  | Other reasons | 6.28 (15) |
| Existing alternative uses of LLINs (n=239) | Ropes for tying and covering items | 20.92 (50) |
|  | Crops and seedlings fencing | 7.53 (18) |
|  | Chicken coop | 5.02 (12) |
|  | Other alternative uses | 7.53 (18) |
|  | No alternative uses | 59 (141) |
| Causes of alternative uses (n=239) | LLIN being worn-out | 12.55 (30) |
|  | LLIN being torn | 23.01 (55) |
|  | Other reasons | 5.44 (13) |
|  | No alternative uses | 59 (141) |

Another social factor for bed net change was seasonal variability (dry and rainfall seasons). The participants stated that mosquito breeding sites reduced drastically during the dry season, while their numbers increased during the rainy season. Therefore, the rainy season is the perpetuator for them to change their bed nets since there is a rapid increase in mosquitoes due to the increasing in number of breeding sites than dry season. Although, there were few participants reported changing their nets was influenced by a need to use in alternative uses.

> *“I normally change the LLIN twice a year, the reason which makes me change LLIN twice is when the rainy season starts since there is increase in mosquito population compare to dry season. You can remain with the bed net regardless is torn during dry season but immediately when rainy season commences, I change the LLIN” (Male, Mlabani). “The reason made me change my LLIN is when the fishing season starts, I see the old net is meaningless so I use it for fishing, therefore I buy the new one for mosquito protection” (Female, Katindiuka).*

### Disposal practices of old LLINs and challenges

More than half of the participants reported to dispose of their bed nets when they are old or worn-out 57.81% (n=384). The most common reported disposal mechanisms were burning 30.73% (n=384), disposing of with other garbage 14.84% (n=384) and alternative uses 8.59% (n=384) and 3.65 others (Table 5). From FGD participants confirmed that the entirely worn-out LLIN should be burned down to ashes because it was useless, therefore it should be burned down along with other household garbage or wastes. The participants explained that they burn the old nets to reduce environmental pollution as they do not disintegrate easily when just left out or buried, as one community member said:

> *“It just like any other wastes you just burn them down; you burn them on the garbage pit or if you do not have a garbage pit you dig it and burn the LLIN with other wastes” (Female, Minepa).*

**Table 5:**
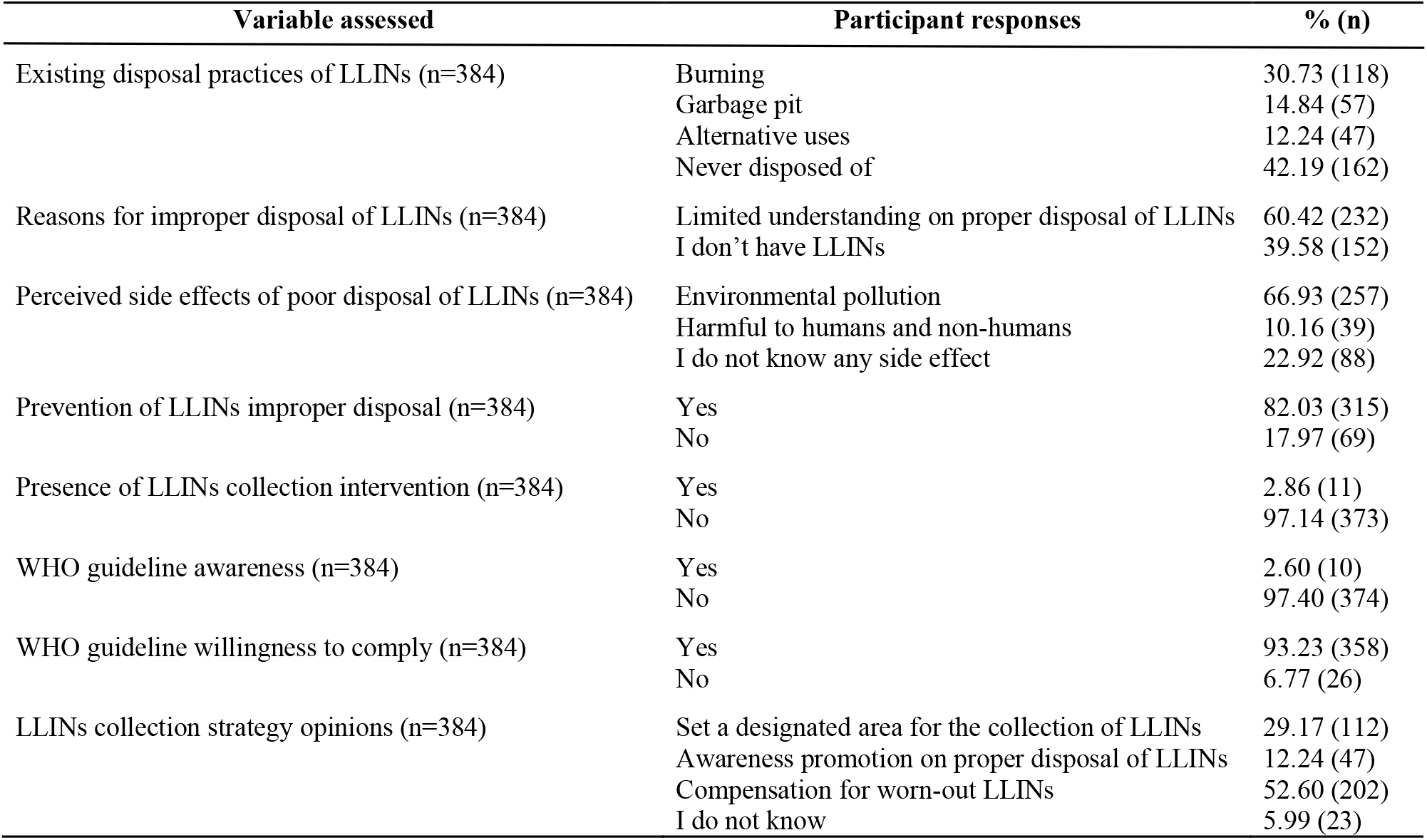
Disposal practices and WHO guideline awareness on the proper disposal of old LLINs

| Variable assessed | Participant responses | % (n) |
| --- | --- | --- |
| Existing disposal practices of LLINs (n=384) | Burning | 30.73 (118) |
|  | Garbage pit | 14.84 (57) |
|  | Alternative uses | 12.24 (47) |
|  | Never disposed of | 42.19 (162) |
| Reasons for improper disposal of LLINs (n=384) | Limited understanding on proper disposal of LLINs | 60.42 (232) |
|  | I don't have LLINs | 39.58 (152) |
| Perceived side effects of poor disposal of LLINs (n=384) | Environmental pollution | 66.93 (257) |
|  | Harmful to humans and non-humans | 10.16 (39) |
|  | I do not know any side effect | 22.92 (88) |
| Prevention of LLINs improper disposal (n=384) | Yes | 82.03 (315) |
|  | No | 17.97 (69) |
| Presence of LLINs collection intervention (n=384) | Yes | 2.86 (11) |
|  | No | 97.14 (373) |
| WHO guideline awareness (n=384) | Yes | 2.60 (10) |
|  | No | 97.40 (374) |
| WHO guideline willingness to comply (n=384) | Yes | 93.23 (358) |
|  | No | 6.77 (26) |
| LLINs collection strategy opinions (n=384) | Set a designated area for the collection of LLINs | 29.17 (112) |
|  | Awareness promotion on proper disposal of LLINs | 12.24 (47) |
|  | Compensation for worn-out LLINs | 52.60 (202) |
|  | I do not know | 5.99 (23) |

There were participants who reported that they often did not know where the old bed nets were kept when they were replaced with new ones. For those that knew reported to store them for future purposes such as using it when there are visitors, passing it on to relatives, selling to other people with alternative uses or keeping it until alternative uses arise as community member mentioned:

> *“For those of us who use LLINs for fishing, when they are destroyed, we do not return them home instead we dump them into the river” (Male, Mlabani).*

### Alternative uses of old nets

During focus discussion, participants reported LLINs that are due for disposal are either as a result of torn or worn-out, therefore can be put to an alternative use as an indication that the useful life of LLINs has ended. In all focus groups, the length of the bed net’s lifespan was defined by its physical condition to fit for the mosquito protection purposes and its socioeconomic values based on the alternative uses. Participants reported several LLINs alternative uses had previously been seen, or practiced in their villages including; garden fencing, chicken coops for protecting chickens from predators, fishing activities and ropes for tying items (Table 2 and Figure 3). While other minimal alternative uses included; curtains, recycled bottle carriers, washing sponge, bathing sponge, floor scrubber and dish dryer. The most dominant LLIN alternative use reported across all focus groups was used for making ropes due to its steady nature for variety of needs including: ropes for building the house/huts, curtains, clotheslines for hanging clothes and ropes for tying different items for example those who sell charcoals sew charcoal bags with LLIN.

> *“There is a time when we use LLIN for building our houses or huts, because the ropes are a little firm, we tear them apart like ropes and then we use them for building houses” (Male, Kivukoni).*

**Figure 3:**
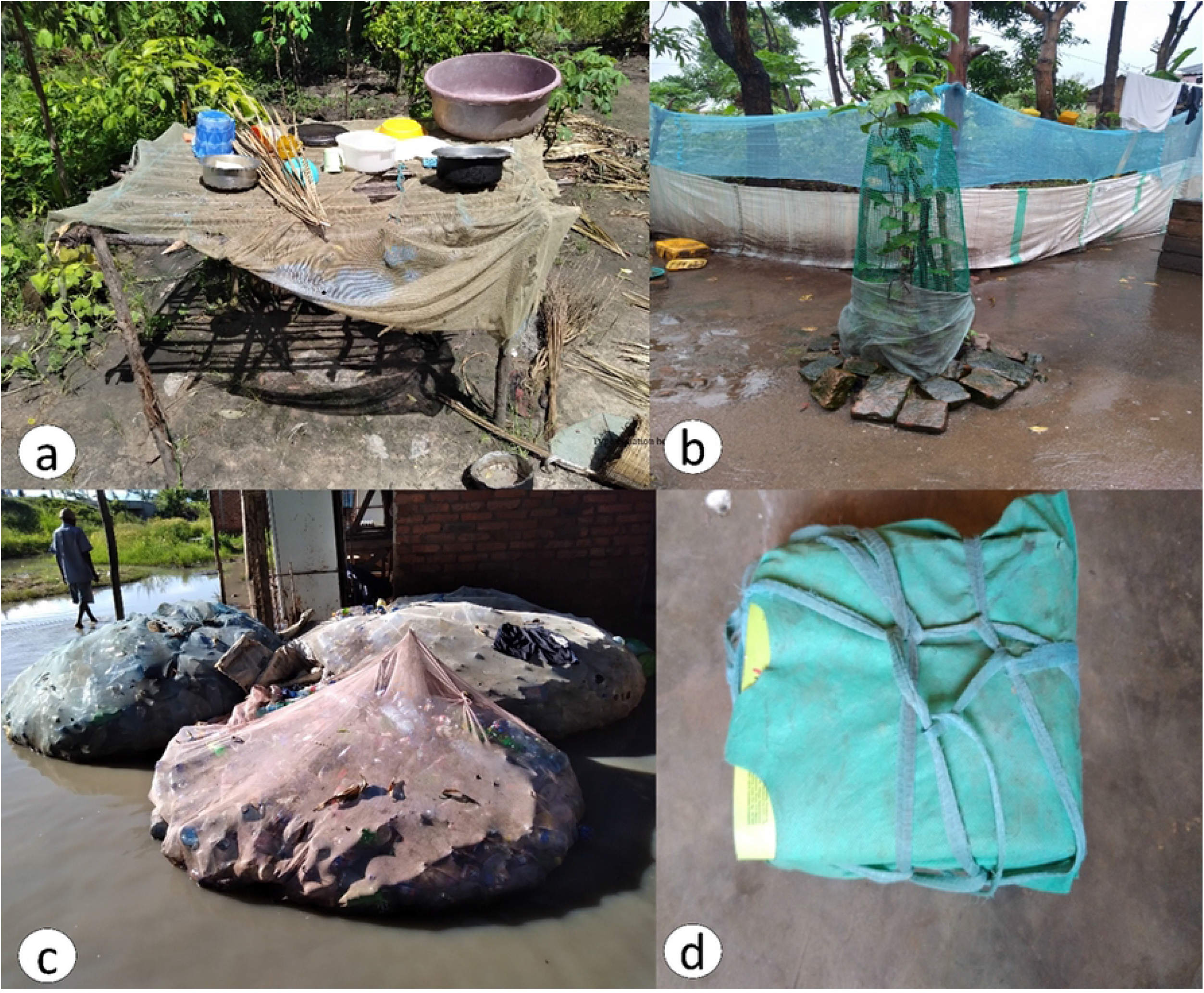
Typical examples of existing alternative use in the study sites, **a**) Illustrates a piece of bed net draped on top of the hut used for drying utensils, **b**) illustrates pieces of bed nets stitched together used for fencing the seedling and garden, **c**) illustrates bed net used as a carrier for the collected recycled plastic bottles and **d**) shows bed net being used as ropes to tie up pupil’s books as one baggage easily to carry

It was further reported that LLINs are used for fencing or as protective barriers for garden around crops and trees seedlings and chicken coops. For the garden fencing is to protect crops or trees seedling from being destroyed by chickens, it is also used to make scary clowns on the rice farms for scaring birds. While on the other hand chicken coop is used to protect chicks from predators such as hawks.

> *“For me when the LLIN is worn-out, I use it for fencing the chick coop at home. Sometime when I grow a garden, I use LLIN to fence the garden to prevent chicken from destroying the crops” (Male, Kivukoni).*

Another dominant alternative use related to fishing, which is an important livelihood activity for communities near the Kilombero river. LLINs were actually preferable to trap small fish which in turn are used as baits to catch larger fish and the practice appears to be justly widespread at Kivukoni village which a located along Kilombero River.

> *“During rainy season like right now, we use LLINs for fishing at the river. About 80% of the residents of this valley use LLINs for fishing because we are residing near along the river valley” (Male, Kivukoni).*

### Improper disposal of LLINs and motives for alternative uses

Major motives for alternative uses of the nets were LLIN being torn 23.01% (n=239), LLIN being worn-out 12.55% (n=239) and other reasons 5.44% (n=239) such as limited understanding of its proper disposal, alternative materials and having more than one net. Although, from FGDs with community members, numerous reasons emerged regarding motives for alternative uses of LLINs. The major motive was LLIN fabric material characteristics considered being steady and durable to fit for other purposes hence saving costs and acting as alternative materials as community member said:

> *“These LLINs which if you touch them are hard, if you look closely, they are the ones that we use for alternative uses especially for fishing, chicken coop and fencing the garden, they are used because of their steadiness” (Male, Mlabani).*

Another most dominant motive for repurposing of the old LLINs alternative uses was poverty, this was echoed by majority of participants across all focus groups. The need for saving costs for working tools such as fishing nets, mesh for fencing to protect chicks and garden instead LLINs were used as alternative materials. For that reason, participants voiced using LLINs as alternative materials for earning to purchase goods and other basic needs such as paying bills, paying for food and school fees. Due to these scenarios, multiple respondents reported turning LLINs for alternative uses.

> *“It is poverty when you find that someone who wants to fish does not have the ability to buy fishing nets, what will he do to earn money, he takes the LLIN and goes fishing to earn money” (Female, Minepa).*

Lack of disposal knowledge and the adverse effects that could be resulted from repurposing or poor disposal of LLINs indicated as the major perpetuator of the poor disposal and alternative uses of the old LLINs. This was audibly echoed by almost all participants to be the major reason for the ongoing alternative uses and poor disposal of the old LLINs. Lack of awareness and knowledge on what to do with the old LLINs were stated to be the motives for alternative uses and poor disposal of the old LLINs by the majority of the participants.

> *The main reason for all this is education, if these people were given the education on proper and improper uses, they would not do that. In this world, even if education is given, the response cannot be 100%, there are people who will go differently, but we will continue to educate each other slowly (Male, Mlabani).*

Another reported reason that perpetuated the alternative uses is having more than one bed net. It was noted that a single household received LLINs from multiple sources for example pregnant woman could get LLIN from clinic, kids from school, mass distribution and ongoing research projects. Some people would tear the LLINs and adjoin them to form a desired size or shape that fit for their purposes. While others with multiples bed nets or their beds were bigger than received bed nets and had no other activities with the bed nets would sell them to those who needs particularly alternative uses.

> *“I have actually witnessed this issue, I have seen it when the aid bed nets were given, you find someone got four, another got five, there are men who were looking for bed nets and buying them to go fishing. People gave out new bed nets and they sold them to people using them for fishing” (Female, Katindiuka).*

### Perceived adverse effects related to poor/improper disposal and alternative uses

Majority of the participants were entirely aware of the harm that poor disposal or repurposing of LLINs could cause the environment and living organisms. They expressed fear of the potential health risks and environmental pollution from the remaining insecticides embedded in the LLINs. Among potential adverse effects captured during focus group discussion were were soil and water pollution, killing of living organisms, air pollution when these LLINs are burned on the open space, mosquitoes developing resistance to the insecticide. The majority of participants expressed their deep fear and concern on the side effects when LLINs used for fishing activities could kill untargeted organisms and premature fish. Consequently, there could be ongoing health risks or disease outbreak related to the insecticides which people are not aware of resulted from consuming fish which are trapped using LLINs.

> *“There are side effects when we dispose of LLINs or go to fish with them, because the LLINs are embedded with insecticides, therefore, we cannot get side effects today or tomorrow. We use them for fishing and the fish could die by insecticide. Consequently, after a certain period of time we could acquire diseases” (Male, Kivukoni).*

However, there were different responses among focus group participants especially males from Kivukoni village who believed the LLINs have no adverse effects since it is perceived to be unfit for mosquito protection. Although, this concern could have been subjective because the majority of men at Kivukoni are involved in fishing activities as a source of income. The participants (males from Kivukoni) further justified that if the embedded insecticide within LLIN could not kill mosquitoes, it shouldn’t pose harm to human beings and fish.

> *“As far as I know, there are no any adverse effects because these LLIN we are told about have insecticides but in their use, we do not see if the mosquitoes actually die. Therefore, we consider these LLINs to be normal in our uses (alternative uses) and they are safe. I believe these bed net which they say have insecticides, but these insecticides are not effective because they are not harmful to the fish. We fish using LLINs and eat those fish, these LLINs themselves do not kill mosquitoes, how will they kill me” (Male, Kivukoni).*
>
> *“I personally did not know if there are any adverse effect on the LLIN because we are given those LLINs to protect ourselves. When I use them to cover myself, I don’t see any harm at times I even touch the LLINs that is why I thought that it does not have any adverse effect even if I use it for other purposes” (Female, Minepa).*

### Community knowledge and awareness of and compliance to WHO guidelines

Out of the 384 respondents of the survey questionnaire, participants 2.6% (n=10) reported that they were aware of the WHO guideline on proper disposal of LLIN, they heard about the guideline predominantly from radio, television and seminar. All interviewed key informants and almost all FGD participants were neither aware of the WHO recommendations on the sound management of the old LLINs nor ever given any instructions on what to do with the old LLINs. FGD participants were entirely not aware of the appropriate methods to dispose of old LLINs and reported neither they had ever given any instructions when bought from shops or received from clinics/schools. Surprisingly, almost all FGD participants were willing to comply with the WHO guideline when the facilitator voiced out the WHO recommendations on the proper disposal of the old LLINs. They also echoed the relevance of the WHO guideline since it protects the environment, public health and other living organisms.

Both KIIs and FGDs participants appreciated the usefulness of the knowledge regarding the WHO recommendations on the proper disposal of the old LLINs. However, the participants argued and emphasized on using the local government (ward/village officers) to disseminate the knowledge to the community members on their regular meetings, this would reinforce the WHO and national environmental management council (NEMC) efforts on the environmental conservation to protect the public health and biodiversity at large.

> *“There is an importance as WHO works with NEMC, what it means that we should do environmental conservation, these are plastics that can cause environmental pollutions and later outbreak of diseases. But also in the river issues, we have those aquatic animals that we use for food. Therefore, when we sustain them and protect them, they become safe for us to consume “ (Male, Mlabani).*
>
> *“These recommendations are good and would be useful as they instructed, maybe we should cooperate with the national malaria control personnel to know the best way to dispose of through burning in high temperature heat in incinerators” (Environmental Officer, Ifakara).*

However, some participants raised concerns on the implementation of the WHO guidelines, some of the recommendations seem to be difficult on the practicality to their settings due to numerous challenges or circumstances which might hinder the collection exercise to be successful. Among mentioned challenges included; lack of enough facilities such as incinerators, strategies to track and collect old LLINs at each household level.

> *“There is a little bit challenge in the recommendations on how to burn those LLINs, I see there they have said that they should be burned in high temperature heat. But now if you look at the places that we should use to burn those LLINs is difficult, the burning facilities of those LLINs are inadequate, for example all Ifakara residents maybe we all burn at St. Francis” (Male, Mlabani).*

Additionally, the participants emphasized on the frequent distribution of LLINs to all community members to ensure accessibility and coverage of LLINs. The participants further argued and proposed the Give and Take (*Nipe Nikupe*) strategy meaning the national malaria control program (NMCP) team should give new LLINs to the community members in exchange to the old ones as WHO guideline suggested to ensure LLIN coverage sustainability but also removal of old LLINs in the community.

> *“Because if you take something from someone and leave something else for him/her, it means that you have not taken something but exchange. Therefore, many people will be convinced, as they will be tired of the old item” (Male, Kivukoni).*

### Recommendation for future: LLINs collection strategies

Focus group discussion participants had different opinions and views regarding on proper collection and disposal of the LLINs. The main collection and disposal strategy of the LLINs proposed by the participants was awareness creation on the adverse effects which could be resulted from the poor disposal of LLINs and alternative uses. It is assumed that the community members are not aware on how to dispose the old/worn-out LLINs on sounding manner as per WHO guideline.

There were notable various opinions on the proposed strategies among men and women since disposing of garbage is more of female chores. Some women participants argued to be given instructions as early as possible at the LLINs delivery stations such as clinics since they are responsible for LLINs caring and disposal. Women therefore emphasized on the awareness creation through medias outreach, formal in-person sessions especially on congregation places such as clinics, schools and village meetings on what to do with the old LLINs. The public health risks and environmental pollution associated with the negative impacts resulted from using of LLINs for alternative use and improper disposal should well be elaborated. This could impact on the awareness raising on the proper disposal of the LLINs as majority of them did not know what to do with the old LLINs.

> *“For example, the WHO can have initiative because we mothers are the ones who are most concerned with these issues, either at the clinic or meetings, then there should be an agenda that talks briefly about how you can store these nets when they are worn out, where they should be sent, so that people know. So, if I have a worn-out LLIN, I hand it over the personnel but there should be often seminars at the clinic where there is large congregation of mothers” (Female, Minepa).*

Focus group discussion participants suggested the NMCP team could train the ward/village leaders and community agents for door-to-door collection strategy every after few weeks/months to collect the old LLINs from households’ level to be disposed of (incinerated) on a sound manner. Furthermore, community members proposed other collection strategies such as replacing the old LLINs with the new one (*“Nipe Nikupe”*) as WHO guideline suggested. They also suggest to be given an incentive to motivate them to collect the LLINs as it is with the recycled plastic bottles.

It was proposed that formulations of rules/regulations and enforcements would improve the outcome. They suggested there should be punishments and fines for non-compliance community members on the proper management of the old LLINs. Consequently, this would lead to the eradication of old/worn-out LLIN wastes in the environment and alternative uses practices.

## Discussion

The use of LLINs for alternative uses rather than disposing them when retired or no longer used for mosquito bites protection is drastically increasing in sub-Saharan Africa including Tanzania. Intensive use of LLINs for alternative uses is influenced by changing decisions and changing frequency of the bed nets. Therefore, the residual insecticides may influence distortion of aquatic habitats when used for fishing and side effects when ingested since some people use for drying fish and washing sponges.

This study shows that bed net replacement was primarily determined by physical condition, torn or worn-out, and the availability of a new bed net. The physical condition included the number and size of holes that allowed mosquitoes to penetrate through [34,35]. The presence of the new bed net was a motivation for replacing the old bed net. This study found that majority of the worn-out and torn bed nets were replaced every 6 months to 2 years. Similarly, other studies reported LLINs to have lifespan of less than three years and that physical condition, age, and the availability of a new bed net are factors for its replacement [34,36–43].

Several alternative uses were documented by this study and the most common across all study sites being observed were; ropes made from LLINs, crops and seedlings protection, chicken coops for protecting chicks against predators, washing and bathing sponges. Ropes were mainly used for tying, covering items, hanging clothes. Mostly in rural areas, it was observed small strips made out of polyester netting were used during as ropes tied with branches and woods on mud houses (huts). Similarly LLINs alternative uses had been documented across other areas in sub-Saharan Africa [9,44–49]. Numerous motives for the bed net alternative uses were revealed from the study; poverty and need for alternative materials being the most common, as LLIN is considered to be a steady fabric material and durable. Another reason was lack of official guidance on how to dispose of old LLINs after it expired or worn-out. Several studies previously reported on the main motives for LLINs alternative uses to be more of economical and available alternative materials and lack of official guidance [5,6,44,50–53]. Furthermore, the findings from this study revealed that majority of the bed nets re-used for alternative uses were worn-out or too torn beyond repair that perceived fading of the insecticides, making the bed net unfit for the protection purpose.

In contrast to other studies on the motives for alternative uses, few community members used LLINs that were still in good condition, they claimed that they were not suitable for mosquito protection due to large meshes (holes), small size of the LLIN in comparison to the printed size, and poor quality of the LLINs. Although a study conducted in Solomon Island had reported on the bed net preference, LLINs was less preferred (especially Olyset^®^ Nets) due to its unfavorable characteristics of large mesh size, easily torn and smaller than the actual size [49]. Sociocultural belief that insecticide-treated bed nets could be the cause of male impotence or infertility, echoed by male participants, was another reason for LLIN alternative. Several other studies have also reported on these concerns and misconceptions about the use of LLINs, particularly the potential risks and impotence (infertility) posed by the insecticides embedded in the bed nets [54–58]. Mutalemwa *et al*., proposed that, in order to dissipate these concerns and misconceptions, all bed net users and recipients should be informed that, when used correctly, the insecticides embedded in LLINs kill and repel mosquitos while remaining safe for humans [9].

As shown on this study, the majority disposal mechanism for LLINs were burning or alternative uses and this is driven mostly due to lack of official guidance on how to dispose of old or worn-out LLINs. This influenced the majority of community members to dispose of their old or worn-out bed nets at their convenience ways. The findings show that the major mosquito control intervention is bed nets particularly LLINs which were freely distributed at antenatal clinics and child immunizations and School Net Program (SNP) and universally distributed by the government through every 3–4 years, with the recent mass distribution in year 2016 [24,43]. These LLINs are made up of polyester and polyethylene materials which are non-degradable plastic and may persist to the environment [13,17,46,59,60]. Despite rapid increase of LLINs distribution coverage and use, there is an increasing environmental and public health concerns when these LLINs are used otherwise or poorly disposed [4].

Besides nets are embedded with insecticides (pyrethroids and pyrroles) which could be toxicant to some biodiversity and the environment, marine organisms can be entangled in LLINs or ingest it depending on the size [61]. Therefore, there should be consideration on biodiversity protection as well as water bodies as stipulated in the SGD goals 6 on clean water and sanitation for all,14 on life below water and 15 on life on land [11]. The widespread alternative uses of LLINs might pose risks to the living organisms for example we observe community members using LLINs for protecting gardens, fishing and washing/bathing sponges so if there are residual when ingested might have posed health risk to human. Further, the residual insecticides might as well as may hinder efforts towards malaria elimination due to the mosquitoes resistance to insecticides and environmental pollution [10,62–64].

Despite the presence of the WHO guideline on handling old LLINs, participants including public and environmental personnel reported not to be aware of the guideline. Although they reported that lack of official guidance and inadequate awareness promotion could be barrier to the guideline propagation. In contrast to previous studies, this study has enlightened on the LLINs disposal practices and its alternative uses and challenges that hinder proper disposal of the old LLINs. Encouragingly, despite being unaware of the WHO guideline on proper disposal of old LLINs, community members were willing to comply with such guidelines. Collection strategies should be agreed upon earlier with community members prior to the net replacement. Evaluation studies conducted in Tanzania and Madagascar on collection of expired LLINs demonstrated that collecting worn-out or retired LLINs may be appropriate if community collection strategies are agreed upon prior to the replacement of new bed nets based on community preferences [9,65]. As a result, the above-mentioned factors should be considered when developing LLINs collection strategy so as to broaden and scale-up the collection of worn-out or retired LLINs.

## Conclusion

This study demonstrate that despite participants’ limited knowledge on WHO guidelines for proper disposal of old LLINs, majority of community members are willing to comply with the guidelines. Since alternative uses sometimes referred to as repurposing of old nets, proper guidelines should be developed ensure that repurposing of old LLINs do not cause harm to human health and the environment. Therefore, there should be a designated point set collecting these nets, although collection strategies should be agreed upon within the community members prior to replacement, as a resident gives the old LLIN and gets the new one (compensation for worn-out nets in terms of money or a new net). Furthermore, comprehensive efforts are therefore necessary to address challenges attributed to alternative uses and poor disposal of old nets as more LLINs get distributed.

## Abbreviations

DSO: Disease Surveillance Officer
EMO: Environmental Management Officer
FGDs: Focus Group Discussions
ITN: Insecticide-treated Bed Nets
KIIs: Key Informant Interviews
LLINs: Long-Lasting Insecticidal Nets
MFP: Malaria Focal Person
NEMC: National Environment Management Council
NMSP: National Malaria Strategic Plan
SNP: School Nets Program
UNEP: United Nations Environment Programme
WHO: World Health Organization

## Author’s contribution

SJS, EWK, WPM and JDN were involved in concept conceptualization and study design. SJS, JDN and WPM were involved in data collection. SJS, YAM, HMM and LLM conducted data analysis. SJS wrote the first draft of the manuscript. EWK, DWL, HMM MFF and WPM reviewed the manuscript. All authors read and approved the final manuscript.

## Authors’ details

1) Ifakara Health Institute, Environmental Health and Ecological Sciences Department, P. O. Box 53 Ifakara, Tanzania. 2) The Nelson Mandela African Institution of Science and Technology, School of Life Sciences and Bio Engineering, P.O Box 447 Arusha, Tanzania. 3) University of Glasgow, Institute of Biodiversity, Animal Health and Comparative Medicine, G12 8QQ, UK. 4) Jaramogi Oginga Odinga University of Science and Technology, Department of Biomedical Sciences, P.O Box 210-40601 Bondo, Kenya. 5) University of the Witwatersrand and the Centre for Emerging Zoonotic and Parasitic Diseases, National Institute for Communicable Diseases, School of Pathology, Faculty of Health Sciences, Johannesburg, South Africa

## Acknowledgment

We would like to thank the district malaria focal persons, vector surveillance officers, environmental and health officers from both Ifakara town council and Ulanga District Council, Ward/Village leaders and community members for their participation in this study. We would also like to extend our sincere gratitude to Ms. Felista S. Tarimo, Ms. Silvina Mjenga, Ms. Jane Chavuka and Ms. Carolina Ndwata for their assistance during data collection. We are also grateful to IHI Ifakara branch team for their guidance, logistical and technical support during the study. We also thank Ms. Edith P. Madumla for their guidance throughout and Ms. Najat Kahamba for drawing a map of the study areas.

## Availability of data and materials

The datasets used and/or analysed during the current study are available from the corresponding author on reasonable request.

